# Improving proteomic dynamic range with Multiple Accumulation Precursor Mass Spectrometry (MAP-MS)

**DOI:** 10.1101/2025.05.14.653938

**Authors:** Teeradon Phlairaharn, Ariana E. Shannon, Xinlei Zeng, Dong-Jiunn Jeffery Truong, Erwin M. Schoof, Zilu Ye, Brian C. Searle

**Affiliations:** Department of Quantitative Health Sciences, Mayo Clinic, Rochester, MN, USA; TUM School of Management, Technical University of Munich, 74076 Heilbronn, Germany; State Key Laboratory of Common Mechanism Research for Major Diseases, Suzhou Institute of Systems Medicine, Chinese Academy of Medical Science & Peking Union Medical College, Suzhou, China; Institute for Synthetic Biomedicine, Helmholtz Munich, Neuherberg, Germany; Department of Biotechnology and Biomedicine, Technical University of Denmark, Lyngby, Denmark

**Keywords:** peptide identification optimization, peptide quantification optimization, orbitrap mass analyzer, multiplexing, mass spectrometry, data-dependent acquisition, data-independent acquisition

## Abstract

Orbitrap (OT) -based mass spectrometer platforms are a gold standard in high-resolution mass spectrometry, where their primary disadvantage is slower-scanning speed in comparison to time-of-flight or linear ion trap mass analyzers. In this study, we utilize long OT transients to extend the precursor dynamic range by modifying the selected ion monitoring method to multiplex several precursor *m/z* ranges from 400 to 1000 *m/z* into a single scan called “<u>M</u>ultiple <u>A</u>ccumulation <u>P</u>recursor <u>M</u>ass <u>S</u>pectrometry” (MAP-MS). Our approach requires no software or hardware modifications and hides the additional ion accumulation steps during the time it takes to make other Orbitrap measurements, producing precursor spectra with nearly 2× dynamic range and essentially no consequences. We collected data using both data-dependent acquisition (DDA) and data-independent acquisition (DIA) methods to evaluate a range of approaches. With DDA, MAP-MS precursor quantification improves with higher quality measurements. At the same time, DIA detection is enhanced by up to 11% when combining precursor and tandem mass spectra for peptide detection.

## INTRODUCTION

Bottom-up proteomics using mass spectrometry (MS) is a powerful technique for global proteome profiling, where instruments typically record two types of scans.^1^ MS1 spectra measure intact peptide *m/z* ratios to provide an overview of the molecular species present in the sample at a given retention time. MS/MS spectra measure peptide fragment ions, offering detailed structural and sequence information from the *m/z* ratios of generated fragments. In DDA, precursors with the highest intensities are targeted for MS/MS analysis,^2^ and quantification is typically performed using MS1 signals. In contrast, DIA^3^ allows for quantification using either MS1 or MS/MS signals,^4^ and identification in bottom-up proteomics typically relies on a combination of MS1 and MS/MS signals.^5^

One challenge in bottom-up proteomics is the dynamic range limitations of mass spectrometers, where high-abundance precursor species can saturate the ion capacity. This creates a bottleneck in the DDA quantification analysis when high-abundance ions saturate (e.g., the C-Trap), preventing low-abundance species from accumulating sufficient copies for detection. Fundamentally, the difference between standard (non-trapping) quadrupole ToF mass spectrometers (QToF-MS) and trapping devices like Orbitraps™ (OT) or linear ion traps (LIT) is how long ions spend in proximity inside the instrument. For QToF-MS, which processes a continuous flow of ions, the ions are constantly moving through the instrument, meaning their interaction with any part of the mass spectrometer is usually less than 1 millisecond (ms). In contrast, traps allow ions to accumulate inside the instrument for a much longer duration before being analyzed, which is fundamentally different from the more transient behavior seen in non-trapping quadrupoles or ToF systems.

With OTs, LITs, and other instruments that trap ions, dynamic range limitations can be significant. Automatic gain control (AGC) and other ion injection regulation systems help manage ion capacity to prevent ion accumulation and mitigate dynamic range limitations that can impact data quality. However, high-abundance species can become dominant in the trap, resulting in a limited dynamic range within a single scan. This is particularly noticeable when there is a large disparity in ion abundances, which can overwhelm the system’s capacity to measure both low and high-abundant species within the same scan, even causing space charging effects and ion coalescence in some Orbitrap spectra. In summary, the fundamental difference between non-trapping systems (quadrupole and ToF) and trapping systems (OT and LIT) lies in how long ions remain inside the instrument, which influences their space charge limitations and the overall ability of the system to handle a wide dynamic range of signals.

Meier et al. established an acquisition method known as ‘BoxCar’ using the Q Exactive™ HF hybrid quadrupole-Orbitrap mass spectrometer to improve intrascan dynamic range at the expense of not collecting MS/MS spectra.^6^ This method lengthens cycle time by filling multiple *m/z* ranges with longer ion injection time (IIT), allowing low-abundance species to be detected in precursor scans and performing accurate mass and time (AMT) analysis on the MS1 level. Newer approaches mix BoxCar measurements with some MS/MS acquisition by decreasing MS1 resolution and ion accumulation time.^7^ When applying this method, finding a balance between reasonable cycle time, the number of MS/MS scans, and the selectivity of precursor scans is crucial.

Here, we introduce a multiplexed acquisition strategy termed “Multiple Accumulation Precursor Mass Spectrometry” (MAP-MS), which extends the dynamic range and quality of MS1 by increasing the efficiency of parallelized transient recording and ion injections to accumulate multiple packets of ions across several precursor mass ranges. MAP-MS borrows conceptually from variable isolation MSX,^8^ but is applied to precursor isolations rather than DIA windows. In this study, we demonstrate that MAP-MS DIA analysis enhances true identification by 11%, based on the analysis of 125 ng of HeLa peptide digest acquired through MAP-MS, when searching in library-free mode. Additionally, quantification in DDA analysis is enhanced, as evidenced by the improved coefficient of variation (CV) values obtained when quantification is performed at the precursor level.

## EXPERIMENTAL SECTION

### Sample preparation and LC mass spectrometry

Pierce HeLa Protein Digest Standard (Thermo Fisher Scientific, catalog number: 88328) was used to perform LC-MSMS analysis. The HeLa protein digest was separated using an Aurora Ultimate TS 25×75 C18 UHPLC column (IonOpticks) in a 1-column setup with a 1-hour gradient and an EASY-nLC 1200 (Thermo Fisher Scientific). We used 125 ng injections for all measurements.

### Standard DDA data acquisition

The scan sequence began with a target selective ion monitoring (tSIM) scan as a precursor ion spectrum on OT at 60k resolution, covering a scan range of 350.4 - 1200.8 *m/z*, a normalized AGC target of either 100% (1e6 ions) or 300% (3e6 ions), and a maximum injection time (IIT) of 24 ms, with the RF lens set to 40%. Precursors for MS/MS analysis were selected in a Top20 fashion, and only charge states 2-6 were included. For MS/MS analysis, precursors were isolated using a 2 *m/z* isolation window and fragmented using HCD with 27% normalized collision energy (NCE). The normalized AGC target was set to 50% (5e4 ions), and the maximum IIT was set to “Auto”.

### Standard DIA data acquisition

The scan sequence began with a tSIM scan as a precursor ion spectrum on OT at 60k resolution, covering a scan range of 390 - 1010 *m/z*, a normalized AGC target of either 100% (1e6 ions) or 300% (3e6 ions), and a maximum IIT of 60 ms, with the RF lens set to 40%. The MS/MS analysis used HCD targeted MS2 (tMS2) measurements (27% NCE), where the precursor isolation range was defined in a mass list table from 400 to 1000 *m/z* with 16 *m/z* staggered windows^9^ to achieve selectivity of 8 *m/z* (**Sup. Tab. S1)**. The normalized AGC target was set to 1000% (1e6 ions), and the maximum IIT to 60 ms. Mass list tables for tSIM and tMS2 in this study were generated using the Encyclopedia software package.^10^

### MAP-MS DDA data acquisition

MAP-MS DDA used the same acquisition strategy as Standard DDA, except with a different MS1 acquisition approach. The MS1 sequence began with six tSIM accumulations measured in a single spectrum configured at 60k resolution across a scan range of 350.4 - 1200.8 *m/z*. The isolation windows were as follows: 126 *m/z* centered at 413.4378 *m/z*, 90 *m/z* centered at 521.4869 *m/z*, 89 *m/z* centered at 611.0276 *m/z*, 100 *m/z* centered at 705.5706 *m/z*, 135.1 *m/z* centered at 823.124 *m/z*, and 310 *m/z* centered at 1045.7252 *m/z* (**Sup. Tab. S2)**. These isolation widths were determined based on the distribution of peptide precursor *m/z*s in the Pan-Human dataset.^11^ The normalized AGC target was 1000% (1e6 ions), with a maximum IIT of 4 ms per window (24 ms total max IIT) and an RF lens configuration at 40%. MS/MS were selected and measured in the same way as for Standard DDA.

### MAP-MS DIA data acquisition

Again, MAP-MS DIA used the same acquisition strategy as Standard DIA, except for a different MS1 approach. Here, the scan sequence began with six tSIM scans as precursor ion spectra on OT at 60k resolution, with a scan range of 390 - 1010 *m/z*. The isolation windows were as follows: isolation window 91 *m/z* centered at 445.9526 *m/z*, 78 *m/z* centered at 530.491 *m/z*, 78 *m/z* centered at 608.5265 *m/z*, 84 *m/z* centered at 689.5634 *m/z*, 106 *m/z* centered at 784.6065 *m/z*, and 163.1 *m/z* centered at 919.1677 *m/z* (**Sup. Tab. S3)**, The normalized AGC target was 1000% (1e6 ions), with a maximum IIT of 10 ms per window (60 ms total max IIT) and an RF lens configuration at 40%. MS/MS were measured in the same way as for standard DIA.

### Chromatogram library generation

Gas-phase fractionation^12,13^ (GPF) was performed to generate a chromatographically matched DIA-based library^10^ for this study. For this, we injected 125 ng HeLa tryptic digest six times covering different 100 *m/z* ranges between 400 - 1000 *m/z* (400 - 500 *m/z*, 500 - 600 *m/z*, 600 - 700 *m/z*, 700 - 800 *m/z*, 800 - 900 *m/z*, and 900 - 1000 *m/z*) using the same LC methods for single shot analysis. For standard DIA, the scan sequence began with a tSIM scan acquired as a precursor ion spectrum on the OT at 60k resolution, using a normalized AGC target of 1000% (1e6 ions), a maximum IIT of 10 ms, an RF lens setting of 40%, and no multiplexed ions. MS/MS analysis parameters were identical to those used in the “Standard DIA acquisition.” The isolation windows were configured as detailed in the Supporting Information (**Sup. Tabs. S5, S7, S9, S11, S13**, and **S15**). For MAP-MS DIA, all parameters for MS and MS/MS analysis were identical to those used in standard DIA for GPF, except that the number of multiplexed ions was set to six. The isolation windows were configured as detailed in the Supporting Information (**Sup. Tabs. S4, S6, S8, S10, S12**, and **S14**).

### Precursor comparison data acquisition

We implemented a single method to characterize multiple types of precursor measurements simultaneously. This method collected seven different types of precursor spectra, each roughly covering 400-1000 *m/z* in succession. The first MS1 approach was modeled after BoxCar,^6^ measuring six isolation windows in six MS1 spectra (36 distinct *m/z* windows, detailed in **Sup. Tab. S16**). Each precursor ion spectrum was acquired using a SIM scan on the Orbitrap at 30k resolution, with a normalized AGC target of 1000%, a maximum IIT of 10 ms, an RF lens setting of 40%, and multiplexed ions enabled to cover distinct mass ranges. MS1 approaches 2, 3, 4, and 5 are different approaches to MAP-MS using alternate windowing schemes with 20 windows (Max IIT=1.25 ms), 12 windows, (Max IIT=2 ms), 6 windows (Max IIT=4 ms), and 3 windows (Max IIT=8 ms), with window schemes detailed in **Sup. Tabs. S3, S17, S18**, and **S19**. The Max IIT settings were chosen to keep the summed ion injection times within a total of 20-25 ms. The sixth MS1 approach used the tSIM configured for 1000% AGC (1e6 ions) while the seventh MS1 approach used the standard MS1 configuration for 100% AGC (1e6 ions). No MS/MS were collected in this acquisition method.

### Bioinformatics

DDA raw files were processed using FragPipe^14^ version 22.0 for all searches and its built-in search engine MSFragger.^15^ For Thermo data, raw files were converted to mzML files by ProteoWizard. Files from different samples were processed separately. The data type was set to “DDA”, and the “default” workflow was used. Files from the same sample were given the same “Experiment” name and “Bioreplicate” number. The Uniprot with “UP000005640_9606” entries released in 2022 was specified as a protein sequence database. IonQuant was used for protein-level quantification, and the MaxLFQ approach was set. All other options were kept as default. Spectronaut (version 19.0.240604.62635) (Biognosys) was used to process DIA data for directDIA via a spectrum-centric approach^16^ and for library-based searching using the GPF library. False-discovery rates (FDRs) for peptide spectral matches (PSMs), peptides, and proteins were set at 1%. Trypsin/P was chosen to ensure enzyme specificity. The peptide length in the search space ranged from seven to fifty-two amino acids, with a maximum allowance of two missed cleavages. Cysteine carbamidomethylation was specified as a fixed modification, while N-terminal acetylation and methionine oxidation were selected as variable modifications. Default settings were applied for all other parameters.

### Statistics

Output tables from FragPipe and Spectronaut were imported into RStudio (Version 2024.04.1). For DDA analysis, all detected peptides and proteins are retained initially. Subsequently, peptides and protein groups with zero intensity values are filtered out. Only peptides and proteins in at least two of three replicates were retained for DIA analysis. Afterward, peptides and protein groups with a quantity of one and/or containing at least one NA value are filtered out.

## RESULTS AND DISCUSSION

### Leveraging Long Orbitrap Transients to Enable MAP-MS

Ion accumulation and ion measurement happen in parallel in OT instruments. However, the time required for those activities varies depending on the type of measurement. With standard OT MS1 measurements, ions are accumulated quickly (typically 1 – 2 ms) across a single, wide precursor isolation range (**Fig. 1A**), but in parallel, 15k resolution DDA measurements take approximately 68 ms (derived from a 64 ms transient and a 4 ms interval between transients), resulting in a scan rate of about 14.7 Hz. As such, while OT instruments parallelize ion accumulation during data acquisition, for MS1 measurements, the instrument typically wastes most of the parallel time.

**Figure 1.**
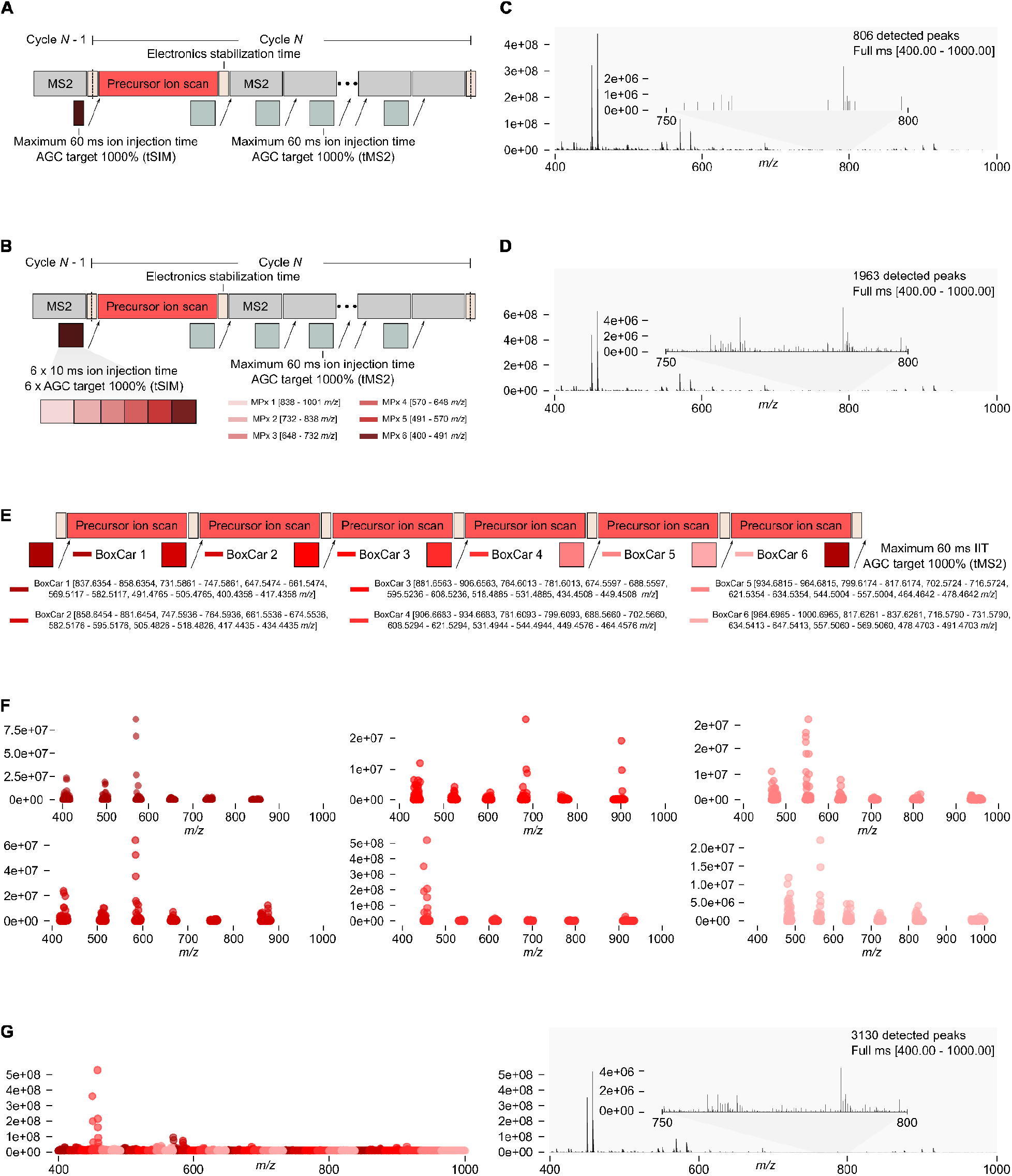
**(A)** Standard MS/MS acquisition scheme. The typical ion accumulation time for the precursor ion scan (MS1) is less than 2 ms despite the available time remaining from the preceding MS/MS scan cycle. **(B)** MAP-MS acquisition scheme using multiplexed tSIMs. The precursor ion scan was acquired in tSIM mode, with the 400–1,000 *m/z* range divided into six subranges (MPx 1 - MPx 6). The maximum IIT for each subrange was set to 10 ms to utilize the remaining time from the preceding MS/MS scan cycle. **(C)** An example precursor scan acquired using the standard acquisition method. **(D)** Matched precursor scan acquired using MAP-MS. In **Figs. 1C** and **1D**, the 750–800 *m/z* region of the full scan is blown up for visualization. With the MAP-MS acquisition method, low-abundance peaks in this region become detectable, whereas they are missed using the standard method due to limitations in dynamic range. In the standard acquisition method, ions from 400–1,000 *m/z* are accumulated together within a fixed IIT. In contrast, MAP-MS divides the range into narrower windows (732–838 *m/z* and 838–1,001 *m/z*), enabling more efficient ion accumulation and enhanced detection of low-abundance signals. **(E)** A 6× MS1 BoxCar acquisition scheme. The schematic illustrates the BoxCar acquisition method applied across six segmented regions spanning 400–1,000 *m/z*. Following the acquisition of BoxCar 1, the adjacent region is sequentially acquired in subsequent scans, continuing this pattern to generate a precursor-only Thermo RAW file. **(F)** Individual peaks acquired in 6× MS1s using the BoxCar method. As a result of the six BoxCar precursor scans, all detectable peaks are shown as individual dots, with intensity on the plot-specific y-axes and the corresponding *m/z* value on the plot-consistent x-axis. The colors in each panel correspond to the defined BoxCar regions shown in **Fig. 1E. (G)** Combined 6× MS1s using the BoxCar method. On the left, the six precursor scans from **Fig. 1F** combined illustrate all detectable peaks using BoxCar scaled to the same y-axis (intensity), where each dot represents a peak’s intensity and its corresponding *m/z* value. On the right, the same data from the combined BoxCar scans are displayed as a spectrum with the same 750–800 *m/z* region blown up for visualization, highlighting the dynamic range similarity between MAP-MS and BoxCar. The spectra in Figures 1C, 1D, 1F, and 1G were acquired immediately adjacent to each other in the same injection to avoid issues with chromatography.

MAP-MS multiplexes precursor ions into multiple precursor *m/z* range packets using a scheme analogous to MSX^8^ (**Fig. 1B**). This approach is similar to BoxCar,^6^ where several ultra-high resolution MS1 spectra are accumulated for very long periods (typically 240 ms) in narrow precursor *m/z* ranges, resulting in high-quality precursor scans. The long MS1 durations mean that there is no time to collect fragmentation data; as such, peptides are identified using accurate mass and time matching (AMT).^17^ In contrast, MAP-MS hides ion accumulation inside the time used to record other OT transients, allowing extra accumulation over a relatively long period. In addition, MAP-MS only acquires a single MS1 per cycle, resulting in essentially no change in the number of acquired spectra and requires no software or hardware modifications, making it essentially consequence-free.

As we present it, MAP-MS is designed to perform six injections for six precursor ion packets across different mass ranges (**Sup. Fig. 1**). To effectively hide MAP-MS ion accumulation, tuning the accumulation time is essential. In a modern high-field Orbitrap, roughly 58 ms are available within the time it takes to make 15k resolution DDA measurements. Since 6 ms are required for electronics stabilization per injection, this leaves 22 ms for collecting ions from all six windows in each MS1. With MAP-MS, we accumulate ions from higher m/z windows first, which helps prevent the loss of high m/z ions that are more weakly confined by the RF pseudopotential in the C-Trap or ion routing multipole (IRM) and are thus more susceptible to space-charging effects. If accumulation were to begin with low m/z ions, these more tightly confined ions could crowd the trap, causing high m/z ions to be lost when the trap becomes overfilled.

To illustrate the effect of MAP-MS in comparison to a standard MS1 scan, we compared paired spectra acquired back-to-back. In the standard acquisition method, the precursor *m/z* range from 400 to 1000 *m/z* is isolated as a single group. High-abundance precursors can fill the C-trap’s capacity, leading to the loss of low-abundance precursors, particularly in the higher *m/z* regions, for example, 750 to 800 *m/z* (**Fig. 1C**). In contrast, MAP-MS fills these regions separately from the high-abundance precursors, producing MS1s with higher dynamic range and detects twice the number of ions with distinct *m/z*s, even though the scanning times for both methods are identical (**Fig. 1D**). To further illustrate the MAP-MS method, we compared MAP-MS to an “ideal” BoxCar-like method collecting six spectra containing six sub-precursor *m/z* ranges (6×6) from 400 to 1000 *m/z* for a total of 36 distinct ion isolations (**Fig. 1E**). When the individual accumulations (**Fig. 1F**) are combined, the resulting spectrum produces a very high-quality MS1 (**Fig. 1G**), confirming the level of detailed signal in the MAP-MS spectrum. While the six spectrum BoxCar approach measures more precursor peaks, the significant reduction in scanning time (a factor of six) gives MAP-MS a notable advantage in overall instrument utilization. Essentially, MAP-MS produces precursor spectra that are partway between BoxCar MS1s and standard MS1s without the additional measurement cost of BoxCar methods.

Using a method that contained only precursor measurements, we directly compared standard MS1 acquisitions to MAP-MS approaches with 1× (a normal tSIM), 3×, 6×, 12×, and 20× windows, as well as the 6×6 BoxCar approach applied to a HeLa proteome. Aside from the “ideal” 6×6 BoxCar benchmark, the other scan types were designed to achieve a maximum IIT of 24-25 ms, matching the 22 ms injection window of 15k resolution DDA MS2s. First, we calculated the total peak count in each precursor ion scan across seven different acquisition setups (**Fig. 2A**). Unsurprisingly, when configured for the same AGC, MS1 and targeted tSIM spectra produced nearly identical results. We observed that all MAP-MS methods measured more precursor peaks than the standard MS1 methods but fewer than 6×6 BoxCar, where MAP-MS with six isolations was the best compromise between the number of isolations and the maximum IIT per isolation (1.9× more ions than standard MS1s). The maximum over the median intensity value is a measure of signal-to-noise, and again, while all MAP-MS approaches produced better spectra than the standard MS1 methods, MAP-MS with six isolations produced the highest signal-to-noise of the single spectrum approaches (**Fig. 2B**). Actual ion injection times show that MAP-MS with 3× ion packets sometimes exits early, MAP-MS with 6× or more ion packets always maximizes the total ion injection time available (**Fig. 2C**). For comparison, the 6×6 BoxCar method required 15× the total ion injection time (spread across 6× spectra) that was available to the single spectrum approaches. Based on our results, we conclude that MAP-MS with 6 ion packets, including the electronic stabilization time, represents an optimized setup that can effectively replace the standard acquisition method.

**Figure 2.**
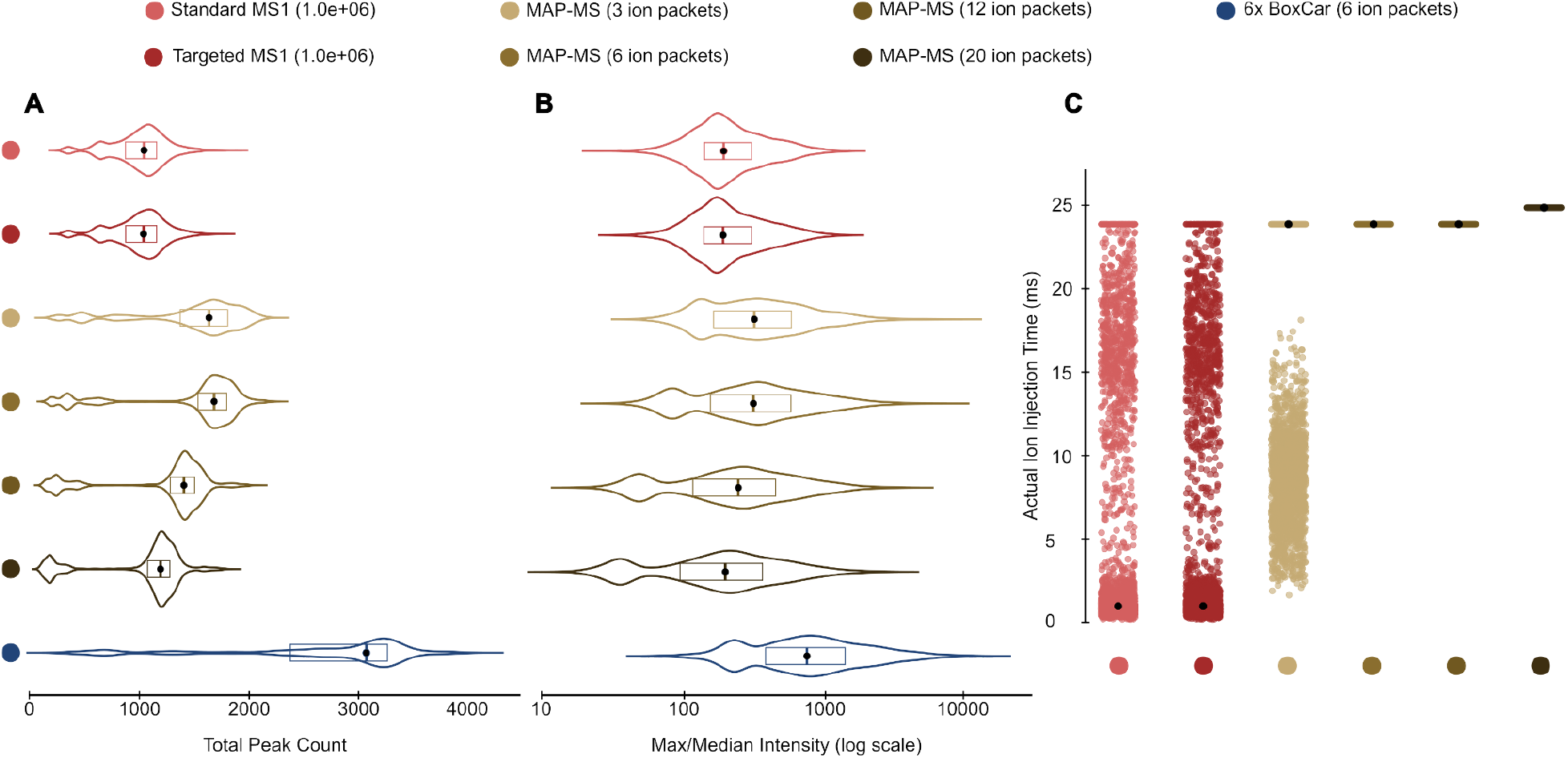
**(A)** The distribution of total peak counts from precursor ion scans obtained using seven different acquisition methods. **(B)** Log-scaled distribution of intensity spread (max/median) from precursor spectra as a measure of the dynamic range. Violin plots indicate the medians as a dot and boxes show the interquartile range. **(C)** Actual ion injection times per precursor spectrum as a measure of accumulation time utilization. Each dot represents the IIT from an individual precursor ion scan for each method, with the median IIT for each method indicated by a black dot.

A critical aspect of MAP-MS is that the AGC target must be applied to each window separately, such that different ion packets get different ion injection times. To demonstrate this, we mapped ion injection time across time for select peaks. We found that the set AGC target dynamically scales the IIT across specific precursor windows in response to the elution of dominant species. Surrounding the example precursor 410.7 *m/z* shown in **Fig. 3**, the total maximum IIT is set to 60 ms, where on average, each window is given 10 ms. However, while the 410.7 m/z peak elutes, the actual IIT for MPx 6 (covering the 400–491 *m/z* range) automatically adjusts to avoid saturating the number of charges and allow the trap to be filled with ions from other windows. This enabled redistribution of IIT to other MPx regions, enhancing the likelihood of detecting low-abundance ions by allowing extended ion accumulation.

**Figure 3.**
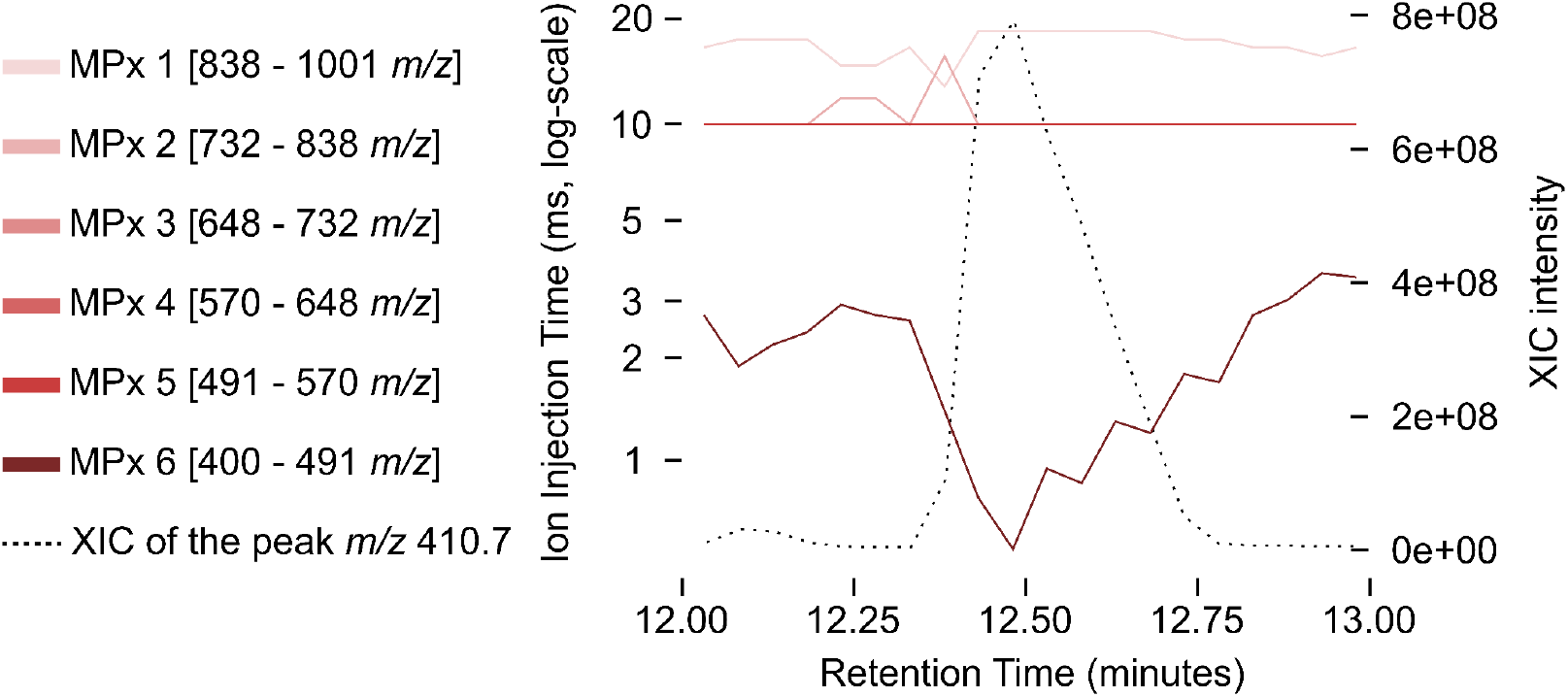
Scaling of the normalized ion injection time during the elution of an example 410.7 *m/z* peak. MPx ion injection times (solid lines) use the left y-axis, while the XIC of peak 410.7 m/z (dashed line) uses the right intensity y-axis. At the time point when the peak at *m/z* 410.7 eluted, the IIT for MPx 6 (covering the 400–491 *m/z* range) was automatically adjusted by the mass spectrometer, allowing the remaining time to be allocated to other regions containing lower-abundance peaks. The left y-axis is in log scale to more clearly visualize the dynamic range changes in IIT.

### Improved DDA through MAP-MS

The increased dynamic range of MAP-MS spectra improves both DDA detection and quantification. To demonstrate this, we analyzed the same HeLa protein digest using identical DDA methods, except we configured precursor scans for either MAP-MS (AGC target = 1e6 charges), standard MS1s with 100% normalized AGC (AGC target = 1e6 charges), or 300% normalized AGC (AGC target = 3e6 charges). While AGC targets above 1e6 can result in increased space charging effects, researchers often configure MS1s to measure ion packets of 3e6 charges to increase sensitivity, albeit at the cost of lowered quantitative accuracy. Without modifications, the MSX AGC target in an Exploris 480 cannot be set above 1e6; therefore, we have not tested if configuring AGC higher with MAP-MS has an advantageous or detrimental effect.

As expected, across triplicate experiments, with MS-Fragger^15^ we detected a total of 75k peptides (5,628 protein groups) with standard MS1s configured to accumulate up to 1e6 charges, and slightly more (76k peptides, 5,747 protein groups) when configured for 3e6 charges (**Fig. 4A**). That improved sensitivity came at the cost of a 32% increase in %CV with IonQuant,^14^ indicating that quantitative reproducibility dropped (**Fig. 4B**). In contrast, when using MAP-MS precursors, we detected nearly 80k peptides (6,134 protein groups) producing a 9% increase in measured proteins (7% over ACG Target=3e6) while simultaneously dropping the quantitative %CV by more than half. Analysis of the detected PSMs revealed that, instead of resampling the same precursors, MAP-MS also increases peptide detection across multiple charge states (**Sup. Tab. 20**), thereby enhancing peptide confidence with new data.

**Figure 4.**
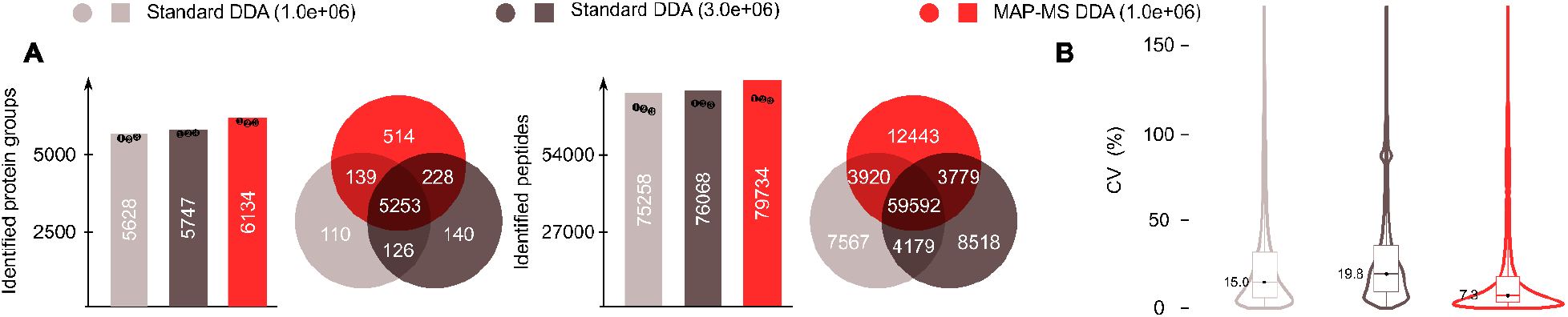
**(A)** Number of identified protein groups (left) and peptides (right) for each acquisition method, visualized using a bar plot where each dot represents the number detected in each of three technical replicates. **(B)** The coefficient of variation (CV) distribution across each acquisition method as violin plots, where the median is indicated as a black dot and the box indicates the interquartile range.

### Enhancing Peptide Detection in DIA through MAP-MS

While DIA is typically less dependent on precursor signals than DDA, in some cases, precursor signals are used for spectrum-centric DIA analysis^18^ and quantification.^4^ Spectrum-centric DIA analysis typically matches the elution patterns of signals in MS/MS to signals in MS1 scans to form pseudospectra.^16,19^ Since MAP-MS provides more accurate and more consistent low-abundance precursor signals, we hypothesized that MAP-MS DIA would improve peptide detection with search engines that utilize this approach. To test this, we used Spectronaut,^20^ a commonly used spectrum-centric DIA approach, to analyze 125 ng injections of a HeLa protein digest standard measured using both standard DIA (with both 1e6 and 3e6 MS1 AGC targets) and MAP-MS DIA. In standard DIA runs using the library-free analysis, we detected a total of 87k peptides (6,611 protein groups) with standard MS1s configured to accumulate up to 1e6 charges, and slightly more than 95k peptides (7,208 protein groups) when configured for 3e6 charges. With MAP-MS DIA, we detected nearly 96k striped peptide sequences (7,531 protein groups), improving overall protein detections by approximately 4% compared to 3e6 AGC Target measurements while maintaining the same quantitative precursor integration accuracy as for DDA.

One potential concern is that these increased detections could also increase the observed peptide False Discovery Rate (FDR). To evaluate this, we first performed an entrapment search,^21^ where, in addition to searching the Homo sapiens database with 4.2M tryptic peptides, and 3.2M tryptic peptides from *C. elegans* were also considered. Since any detections of *C. elegans* peptides are false, we used this to assess the true rate of false discovery outside of Spectronaut (**Fig. 5B**). We found that even with higher reported peptides, MAP-MS DIA falsely identified fewer entrapment peptides, indicating that the increase in detections came without a meaningful penalty.

**Figure 5.**
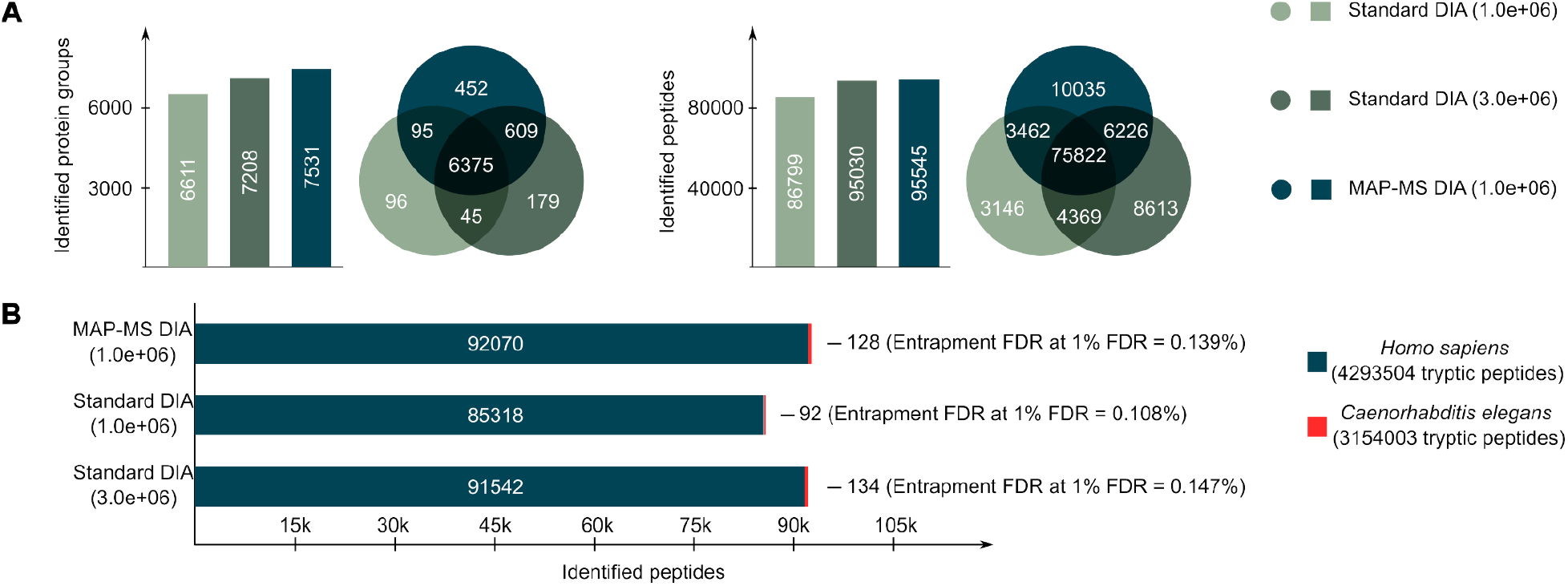
**(A)** The number of identified protein groups (left) and peptides (right) for each acquisition method. Venn diagrams illustrate the overlap of protein groups and peptides identified by the three methods, with shared areas indicating those detected by multiple methods. **(B)** The number of unique human peptide sequences when searching MAP-MS DIA and standard DIA datasets using entrapment, where both *Homo sapiens* and *C. elegans* databases were considered. The number of misidentified entrapment peptides (red) and the percentage of entrapment false discoveries are listed.

MAP-MS can also be applied to gas-phase fractionation, an effective tool for DIA library generation.^22^ In our study, we employed the chromatogram library generation approach,^10^ which generates libraries by analyzing a sample pool using six gas-phase fractions, each configured to analyze a different 100 *m/z* range (**Fig. 6A**). In the MAP-MS scheme, we adjust the MS1 measurement to inject ions from six separate 17 *m/z* windows. For example, the 400-500 m/z GPF-DIA MS1 analyzes the following ion packets: 399.5 – 416.5 *m/z*, 416.5 – 433.5 *m/z*, 433.5 – 450.5 *m/z*, 450.5 – 467.5 *m/z*, 467.5 – 484.5 *m/z*, and 484.5 – 501.5 *m/z*, including a small margin at the top end to capture the entire isotopic packet for each peptide in this range. As a result, MAP-MS DIA showed enhanced performance by detecting 160,076 stripped peptides across six gas-phase fractions, compared to 153,631 peptides detected with the standard DIA method. This 4.1% increase in library size was associated with improved peptide detection sensitivity.

**Figure 6.**
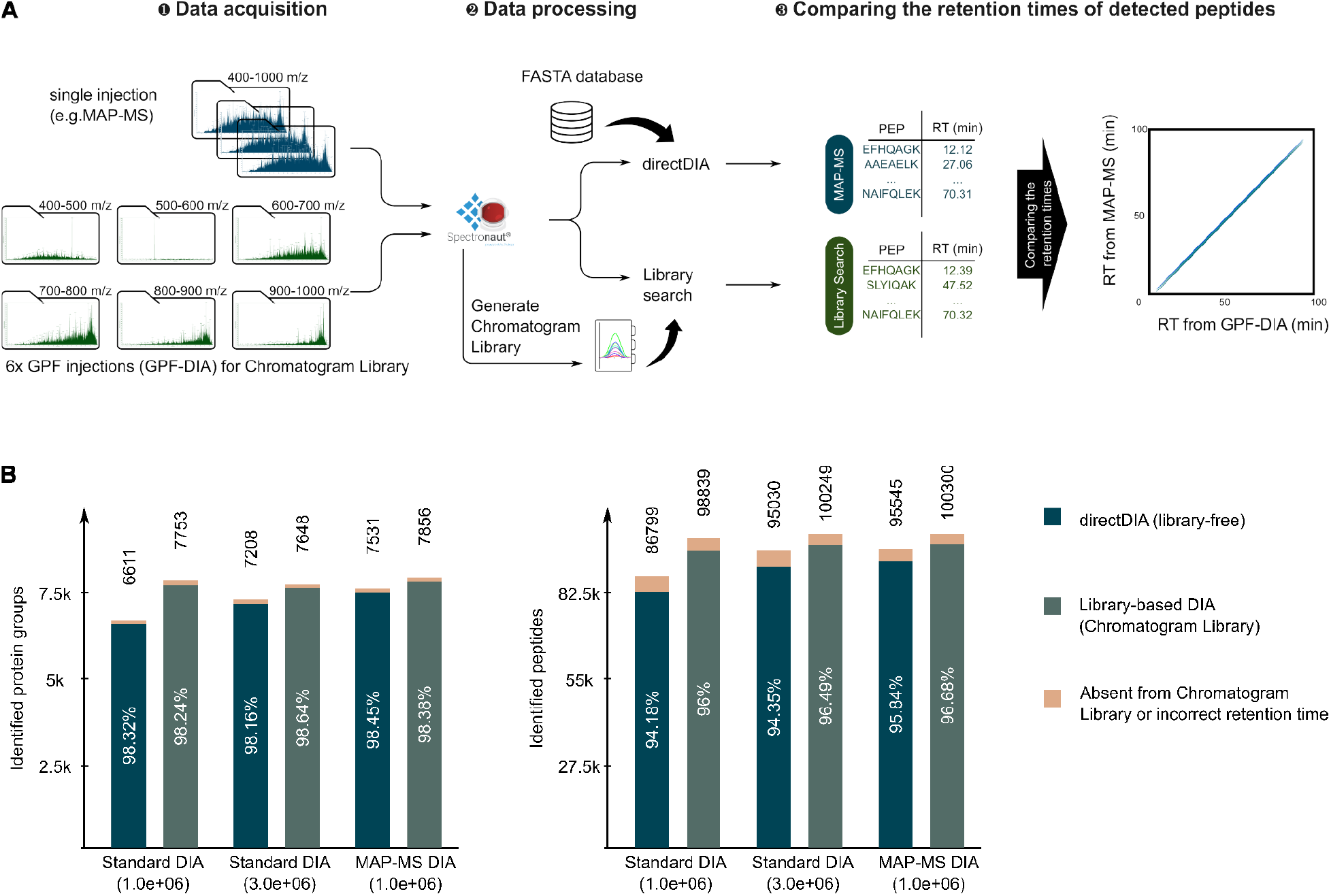
**(A)** Schematic workflow of data acquisition and analysis. Standard DIA and MAP-MS DIA files were analyzed using either directDIA or library searching using the chromatogram library workflow. **(B)** Number of identified protein groups (left) and peptides (right) across acquisition methods after applying RT-based filtering. Identifications that were absent from the GPF-DIA reference or with retention time deviations greater than 90 seconds were marked as incorrect.

We found that, unsurprisingly, larger libraries help identify more peptides. With library searching, we detected 100,300 peptides and 7,856 protein groups using a MAP-MS DIA, representing a 1-3% increase over standard DIA experiments with normal chromatogram libraries (**Fig. 6B**). Additionally, using chromatogram libraries consistently increased detection levels compared to directDIA. While using MAP-MS did not improve the number of detected proteins as dramatically as with DDA, we did observe a consistent increase in the number of true positive peptides that match observed retention times within 90 seconds (**Sup. Fig. 2**). This approach represents an orthogonal method to entrapment for validating the true false discovery rate.

## CONCLUSIONS

MAP-MS is a method to extend the dynamic range of precursor spectra, enabling deeper detection rates with both DDA and DIA as well as significantly more consistent precursor quantification. Since MAP-MS ion accumulation hides within the time it takes to acquire other scans, the approach is essentially “regret-free” and easily implementable on any current generation Thermo Orbitrap mass spectrometer with standard method development options. The approach can drop into any Orbitrap acquisition scheme (including precursor-only approaches), making it applicable to a wide range of analytes beyond peptides.

While we demonstrate MAP-MS in standard proteomics experiments, increasing precursor dynamic range should have significant effects in datasets that span multiple organisms such as metaproteomics^23^ and host/pathogen experiments.^24^ Analysis of high dynamic range biofluids^25^ such as plasma, serum, and urine, should also strongly benefit from improved precursor measurements, especially for DDA. In addition, we believe MAP-MS will particularly help when match-between-runs signal inference is required for a large number of measurements, for example with single-cell proteomics.^26^ Most importantly, we believe that MAP-MS shows a roadmap of how to make more sophisticated use of the ion beam during acquisition using existing hardware and software tools.

## ASSOCIATED CONTENT

### Data availability

This study’s raw mass spectrometry data have been deposited with the ProteomeXchange Consortium via the MassIVE repository. Project accession: PXD054037.

### Supporting information

Ion accumulations and windowing for standard DIA and MAP-MS DIA analysis; Retention time correlation between peptides identified directDIA using MAP-MS DIA and those identified through 6× GPF acquired with standard DIA methods; acquisition parameters for this study.

## AUTHOR INFORMATION

### Corresponding Author

**Brian C. Searle** - Department of Quantitative Health Sciences, Mayo Clinic, Rochester, MN, USA;

### Author Contributions

B.C.S. and T.P. designed the study. B.C.S. and A.E.S. performed the experiments. T.P., X.Z., Z.Y., and B.C.S. performed the data analysis. T.P., D.J.J.T., E.M.S., Z.Y., and B.C.S. contributed input to the method design and data evaluation. T.P. and B.C.S. drafted and revised the manuscript, which has been read and approved by all authors. B.C.S. supervised the work.

## ACKNOWLEDGMENTS

T.P. would like to express his deepest appreciation to H. I. Stewart, D. Chernyshev, C. Hock, E. Denisov, J.M. Garland, J. Petzoldt, A.C. Peterson, B. Hagedorn, Y. Müller, and A. Stefes at Thermo Fisher Scientific, Bremen for great mentorship. T.P. would like to express heartfelt gratitude to T. Bantersons at Roskilde University for the support and motivating influence throughout this journey.

## ABBREVIATIONS

AGC: Automatic Gain Control
DDA: data-dependent acquisition
DIA: data-independent acquisition
GPF: gas-phase fractionation
IIT: ions injection time
LC: liquid chromatography
MAP-MS: Multiple Accumulation Precursor Mass Spectrometry
MS: mass spectrometry
ms: millisecond
OHPF: off-line moderately high-pH reversed-phase fractionation
OT: orbitrap
RT: retention time
tMS2: target MS2
tSIM: target selective ion monitoring;

