## Supporting information for "Improving proteomic dynamic range with Multiple Accumulation Precursor Mass Spectrometry (MAP-MS)"

### Supplementary Figures

**A**

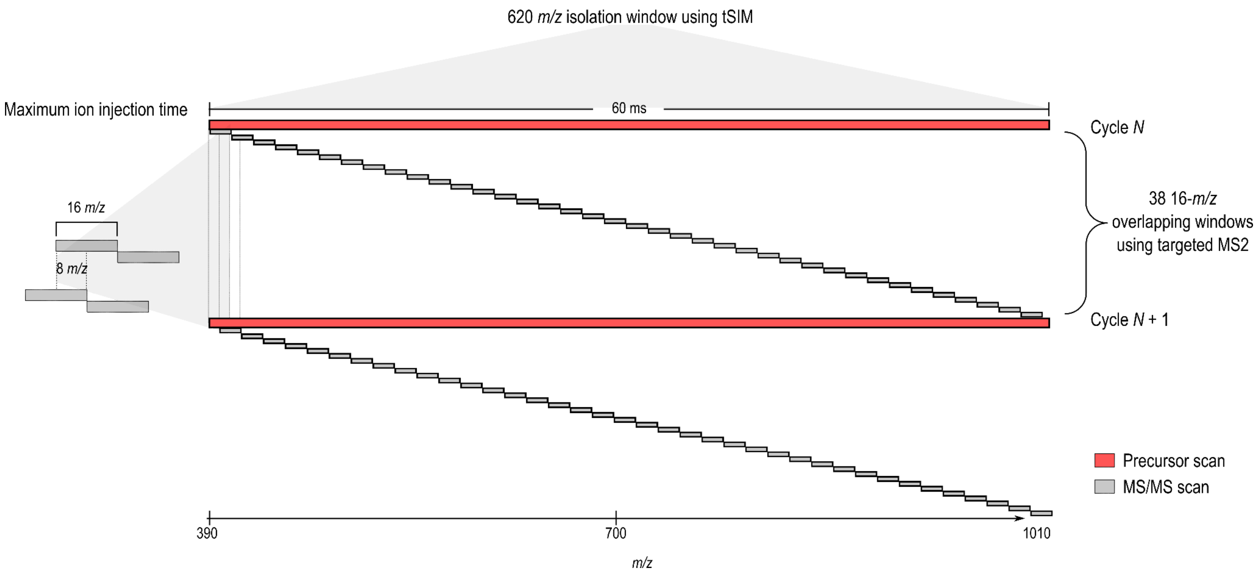


**B**

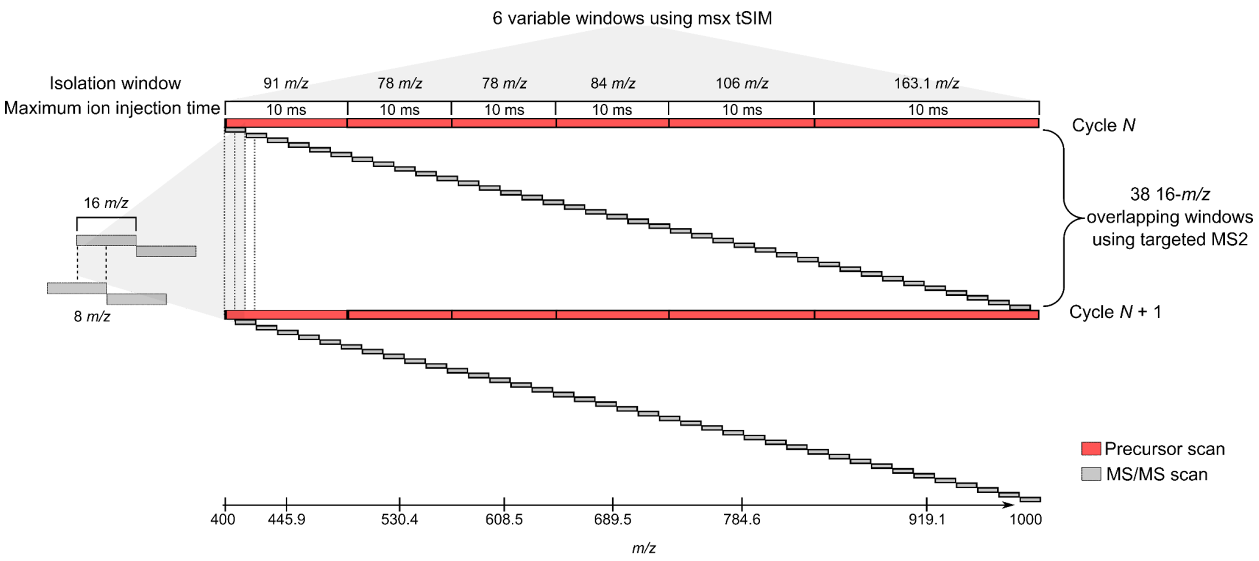


**Supplementary Figure S1: Ion accumulation and windowing for DIA schemes.** (A) Standard DIA acquisition scheme. (B) MAP-MS DIA acquisition scheme. The only change between these methods is the windows and maximum ion accumulation times in the precursor.

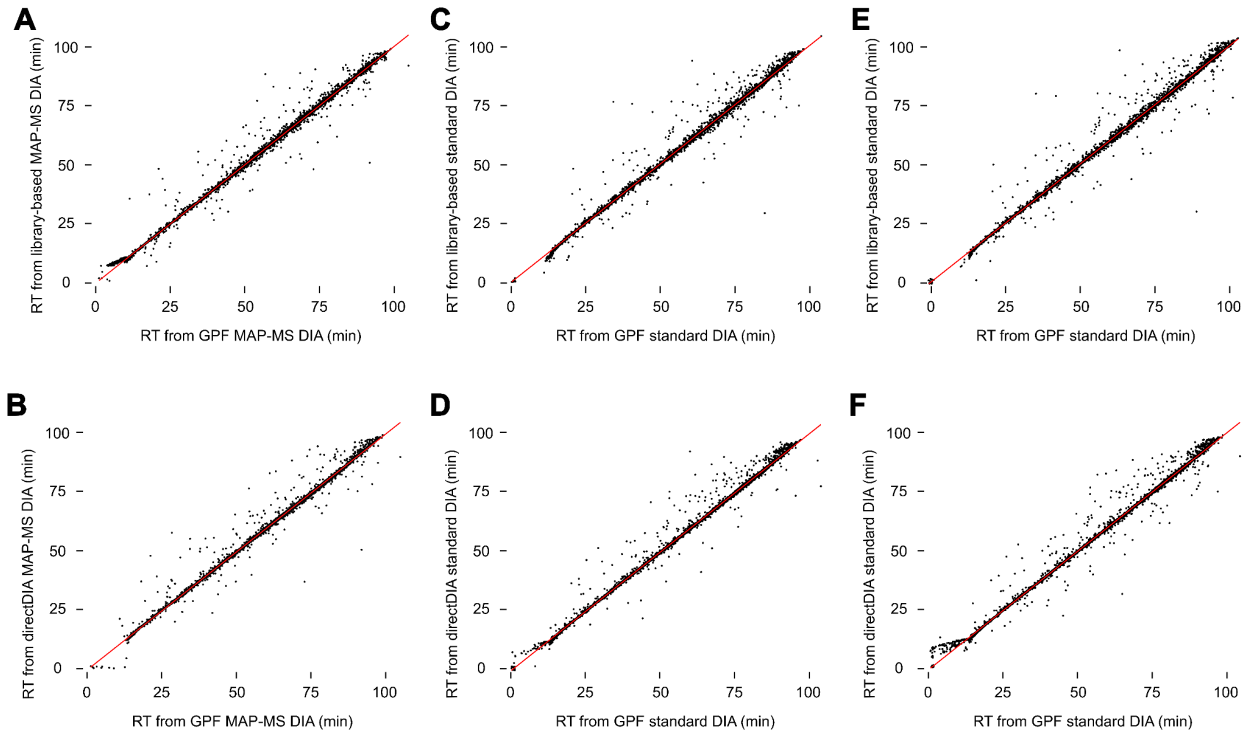


**Supplementary Figure S2: Confirmation of MAP-MS precursors by comparison to GPF-DIA.**

Retention time correlation between peptides identified directDIA using MAP-MS DIA and those identified through 6× GPF acquired with standard DIA methods.

### Supplementary Tables

**Supplementary Table S1: DIA precursor isolation mass list tables for MS/MS spectra (tMS2) (fixed 16 *m/z* isolation windows)**

| Compound | Formula | Adduct | Center mass (*m/z*) | z |
|---|---|---|---|---|
| | | no adduct | 408.4356 | 3 |
| | | no adduct | 424.4428 | 3 |
| | | no adduct | 440.4501 | 3 |
| | | no adduct | 456.4574 | 3 |
| | | no adduct | 472.4647 | 3 |
| | | no adduct | 488.4719 | 3 |
| | | no adduct | 504.4792 | 3 |
| | | no adduct | 520.4865 | 3 |
| | | no adduct | 536.4938 | 3 |
| | | no adduct | 552.5010 | 3 |
| | | no adduct | 568.5083 | 3 |
| | | no adduct | 584.5156 | 3 |
| | | no adduct | 600.5228 | 3 |
| | | no adduct | 616.5302 | 3 |
| | | no adduct | 632.5374 | 3 |
| | | no adduct | 648.5447 | 3 |
| | | no adduct | 664.5520 | 3 |
| | | no adduct | 680.5592 | 3 |
| | | no adduct | 696.5665 | 3 |
| | | no adduct | 712.5737 | 3 |
| | | no adduct | 728.5811 | 3 |
| | | no adduct | 744.5884 | 3 |
| | | no adduct | 760.5956 | 3 |
| | | no adduct | 776.6029 | 3 |

| | | | | |
|---|---|---|---|---|
| | | no adduct | 792.6101 | 3 |
| | | no adduct | 808.6174 | 3 |
| | | no adduct | 824.6248 | 3 |
| | | no adduct | 840.6320 | 3 |
| | | no adduct | 856.6393 | 3 |
| | | no adduct | 872.6466 | 3 |
| | | no adduct | 888.6538 | 3 |
| | | no adduct | 904.6611 | 3 |
| | | no adduct | 920.6683 | 3 |
| | | no adduct | 936.6757 | 3 |
| | | no adduct | 952.6829 | 3 |
| | | no adduct | 968.6902 | 3 |
| | | no adduct | 984.6975 | 3 |
| | | no adduct | 1000.7047 | 3 |
| | | no adduct | 400.4319 | 3 |
| | | no adduct | 416.4392 | 3 |
| | | no adduct | 432.4465 | 3 |
| | | no adduct | 448.4537 | 3 |
| | | no adduct | 464.4610 | 3 |
| | | no adduct | 480.4683 | 3 |
| | | no adduct | 496.4756 | 3 |
| | | no adduct | 512.4828 | 3 |
| | | no adduct | 528.4901 | 3 |
| | | no adduct | 544.4974 | 3 |
| | | no adduct | 560.5046 | 3 |
| | | no adduct | 576.5120 | 3 |
| | | no adduct | 592.5192 | 3 |
| | | no adduct | 608.5265 | 3 |

| | | | | |
|---|---|---|---|---|
| | | no adduct | 624.5338 | 3 |
| | | no adduct | 640.5410 | 3 |
| | | no adduct | 656.5483 | 3 |
| | | no adduct | 672.5555 | 3 |
| | | no adduct | 688.5629 | 3 |
| | | no adduct | 704.5702 | 3 |
| | | no adduct | 720.5774 | 3 |
| | | no adduct | 736.8547 | 3 |
| | | no adduct | 752.5919 | 3 |
| | | no adduct | 768.5992 | 3 |
| | | no adduct | 784.6066 | 3 |
| | | no adduct | 800.6138 | 3 |
| | | no adduct | 816.6211 | 3 |
| | | no adduct | 832.6284 | 3 |
| | | no adduct | 848.6356 | 3 |
| | | no adduct | 864.6429 | 3 |
| | | no adduct | 880.6502 | 3 |
| | | no adduct | 896.6575 | 3 |
| | | no adduct | 912.6647 | 3 |
| | | no adduct | 928.6720 | 3 |
| | | no adduct | 944.6793 | 3 |
| | | no adduct | 960.6865 | 3 |
| | | no adduct | 976.6939 | 3 |
| | | no adduct | 992.7011 | 3 |

**Supplementary Table S2: Precursor mass list tables for MAP-MS DDA (350.4 - 1200.8 *m/z*)**

| Compound | Formula | Adduct | Center mass (*m/z*) | z | Isolation window (*m/z*) |
|---|---|---|---|---|---|
| Boxcar 1 | | no adduct | 413.4378 | 1 | 126 |
| Boxcar 2 | | no adduct | 521.4869 | 1 | 90 |
| Boxcar 3 | | no adduct | 611.0276 | 1 | 89 |
| Boxcar 4 | | no adduct | 705.5706 | 1 | 100 |
| Boxcar 5 | | no adduct | 823.1240 | 1 | 135.1 |
| Boxcar 6 | | no adduct | 1045.7252 | 1 | 310.1 |

**Supplementary Table S3: Precursor mass list tables for MAP-MS DIA (390 - 1010 *m/z*)**

| Compound | Formula | Adduct | Center mass (*m/z*) | z | Isolation window (*m/z*) |
|---|---|---|---|---|---|
| Boxcar 1 | | no adduct | 445.9526 | 1 | 91 |
| Boxcar 2 | | no adduct | 530.4910 | 1 | 78 |
| Boxcar 3 | | no adduct | 608.5265 | 1 | 78 |
| Boxcar 4 | | no adduct | 689.5634 | 1 | 84 |
| Boxcar 5 | | no adduct | 784.6065 | 1 | 106 |
| Boxcar 6 | | no adduct | 919.1677 | 1 | 163.1 |

**Supplementary Table S4: Mass list tables for precursor ion scan (tSIM) between 400 - 500 *m/z* for MAP-MS GPF-DIA**

| Compound | Formula | Adduct | Center mass (*m/z*) | z | Isolation window (*m/z*) |
|---|---|---|---|---|---|
| Boxcar 1 | | no adduct | 407.9353 | 1 | 17 |
| Boxcar 2 | | no adduct | 424.9431 | 1 | 17 |
| Boxcar 3 | | no adduct | 441.9508 | 1 | 17 |
| Boxcar 4 | | no adduct | 458.9585 | 1 | 17 |
| Boxcar 5 | | no adduct | 475.9663 | 1 | 17 |
| Boxcar 6 | | no adduct | 492.974 | 1 | 17 |

**Supplementary Table S5: Mass list tables for MS/MS scan (tMS2) between 400 - 500 *m/z* for MAP-MS GPF-DIA and Standard GPF-DIA (fixed 4 *m/z* isolation windows)**

| Compound | Formula | Adduct | *m/z* | z |
|---|---|---|---|---|
| | | no adduct | 402.4328 | 3 |
| | | no adduct | 406.4346 | 3 |
| | | no adduct | 410.4365 | 3 |
| | | no adduct | 414.4383 | 3 |
| | | no adduct | 418.4401 | 3 |
| | | no adduct | 422.4419 | 3 |
| | | no adduct | 426.4437 | 3 |
| | | no adduct | 430.4456 | 3 |
| | | no adduct | 434.4474 | 3 |
| | | no adduct | 438.4492 | 3 |
| | | no adduct | 442.451 | 3 |
| | | no adduct | 446.4528 | 3 |
| | | no adduct | 450.4547 | 3 |

| | | | | |
|---|---|---|---|---|
| | | no adduct | 454.4565 | 3 |
| | | no adduct | 458.4583 | 3 |
| | | no adduct | 462.4601 | 3 |
| | | no adduct | 466.4619 | 3 |
| | | no adduct | 470.4638 | 3 |
| | | no adduct | 474.4656 | 3 |
| | | no adduct | 478.4674 | 3 |
| | | no adduct | 482.4692 | 3 |
| | | no adduct | 486.471 | 3 |
| | | no adduct | 490.4728 | 3 |
| | | no adduct | 494.4746 | 3 |
| | | no adduct | 498.4765 | 3 |

**Supplementary Table S6: Mass list tables for precursor ion scan (tSIM) between 500 - 600 *m/z* for MAP-MS GPF-DIA**

| Compound | Formula | Adduct | Center mass (*m/z*) | z | Isolation window (*m/z*) |
|---|---|---|---|---|---|
| Boxcar 1 | | no adduct | 507.9808 | 1 | 17 |
| Boxcar 2 | | no adduct | 524.9885 | 1 | 17 |
| Boxcar 3 | | no adduct | 541.9962 | 1 | 17 |
| Boxcar 4 | | no adduct | 559.004 | 1 | 17 |
| Boxcar 5 | | no adduct | 576.0117 | 1 | 17 |
| Boxcar 6 | | no adduct | 593.0194 | 1 | 17 |

**Supplementary Table S7: Mass list tables for MS/MS scan (tMS2) between 500 - 600 *m/z* for MAP-MS GPF-DIA and standard GPF-DIA (fixed 4 *m/z* isolation windows)**

| Compound | Formula | Adduct | *m/z* | z |
|---|---|---|---|---|
| | | no adduct | 502.4783 | 3 |
| | | no adduct | 506.4801 | 3 |
| | | no adduct | 510.4819 | 3 |
| | | no adduct | 514.4838 | 3 |
| | | no adduct | 518.4856 | 3 |
| | | no adduct | 522.4874 | 3 |
| | | no adduct | 526.4892 | 3 |
| | | no adduct | 530.491 | 3 |
| | | no adduct | 534.4929 | 3 |
| | | no adduct | 538.4946 | 3 |
| | | no adduct | 542.4965 | 3 |
| | | no adduct | 546.4983 | 3 |
| | | no adduct | 550.5001 | 3 |
| | | no adduct | 554.502 | 3 |
| | | no adduct | 558.5038 | 3 |
| | | no adduct | 562.5056 | 3 |
| | | no adduct | 566.5074 | 3 |
| | | no adduct | 570.5092 | 3 |
| | | no adduct | 574.5111 | 3 |
| | | no adduct | 578.5128 | 3 |
| | | no adduct | 582.5147 | 3 |
| | | no adduct | 586.5165 | 3 |
| | | no adduct | 590.5183 | 3 |
| | | no adduct | 594.5201 | 3 |
| | | no adduct | 598.522 | 3 |

**Supplementary Table S8: Mass list tables for precursor ion scan (tSIM) between 600 - 700 *m/z* for MAP-MS GPF-DIA**

tSIM scan mass list table for MAP-MS DIA precursor ion scan analysis

| Compound | Formula | Adduct | Center mass (*m/z*) | z | Isolation window (*m/z*) |
|---|---|---|---|---|---|
| Boxcar 1 | | no adduct | 608.0263 | 1 | 17 |
| Boxcar 2 | | no adduct | 625.034 | 1 | 17 |
| Boxcar 3 | | no adduct | 642.0418 | 1 | 17 |
| Boxcar 4 | | no adduct | 659.0494 | 1 | 17 |
| Boxcar 5 | | no adduct | 676.0572 | 1 | 17 |
| Boxcar 6 | | no adduct | 693.0649 | 1 | 17 |

**Supplementary Table S9: Mass list tables for MS/MS scan (tMS2) between 600 - 700 *m/z* for MAP-MS GPF-DIA and standard GPF-DIA (fixed 4 *m/z* isolation windows)**

tMS2 scan mass list table for standard DIA and MAP-MS DIA MS/MS analysis

| Compound | Formula | Adduct | *m/z* | z |
|---|---|---|---|---|
| | | no adduct | 602.5237 | 3 |
| | | no adduct | 606.5256 | 3 |
| | | no adduct | 610.5274 | 3 |
| | | no adduct | 614.5292 | 3 |
| | | no adduct | 618.531 | 3 |
| | | no adduct | 622.5328 | 3 |
| | | no adduct | 626.5347 | 3 |
| | | no adduct | 630.5365 | 3 |
| | | no adduct | 634.5383 | 3 |
| | | no adduct | 638.5402 | 3 |
| | | no adduct | 642.5419 | 3 |

| | | | | |
|---|---|---|---|---|
| | | no adduct | 646.5438 | 3 |
| | | no adduct | 650.5456 | 3 |
| | | no adduct | 654.5474 | 3 |
| | | no adduct | 658.6492 | 3 |
| | | no adduct | 662.551 | 3 |
| | | no adduct | 666.5529 | 3 |
| | | no adduct | 670.5547 | 3 |
| | | no adduct | 674.5565 | 3 |
| | | no adduct | 678.5584 | 3 |
| | | no adduct | 682.562 | 3 |
| | | no adduct | 686.562 | 3 |
| | | no adduct | 690.5638 | 3 |
| | | no adduct | 694.5656 | 3 |
| | | no adduct | 698.5674 | 3 |

**Supplementary Table S10: Mass list tables for precursor ion scan (tSIM) between 700 - 800 *m/z* for MAP-MS GPF-DIA**

| Compound | Formula | Adduct | Center mass (*m/z*) | z | Isolation window (*m/z*) |
|---|---|---|---|---|---|
| Boxcar 1 | | no adduct | 708.0718 | 1 | 17 |
| Boxcar 2 | | no adduct | 725.0795 | 1 | 17 |
| Boxcar 3 | | no adduct | 742.0872 | 1 | 17 |
| Boxcar 4 | | no adduct | 759.095 | 1 | 17 |
| Boxcar 5 | | no adduct | 776.102 | 1 | 17 |
| Boxcar 6 | | no adduct | 793.1104 | 1 | 17 |

**Supplementary Table S11: Mass list tables for MS/MS scan (tMS2) between 700 - 800 *m/z* for MAP-MS GPF-DIA and standard GPF-DIA (fixed 4 *m/z* isolation windows)**

| Compound | Formula | Adduct | *m/z* | z |
|---|---|---|---|---|
| | | no adduct | 702.5692 | 3 |
| | | no adduct | 706.571 | 3 |
| | | no adduct | 710.5729 | 3 |
| | | no adduct | 714.5747 | 3 |
| | | no adduct | 718.5765 | 3 |
| | | no adduct | 722.5783 | 3 |
| | | no adduct | 726.5801 | 3 |
| | | no adduct | 730.582 | 3 |
| | | no adduct | 734.5838 | 3 |
| | | no adduct | 738.5856 | 3 |
| | | no adduct | 742.5874 | 3 |
| | | no adduct | 746.5892 | 3 |
| | | no adduct | 750.5911 | 3 |
| | | no adduct | 754.5929 | 3 |
| | | no adduct | 758.5947 | 3 |
| | | no adduct | 762.5965 | 3 |
| | | no adduct | 766.5983 | 3 |
| | | no adduct | 770.6002 | 3 |
| | | no adduct | 774.602 | 3 |
| | | no adduct | 778.6038 | 3 |
| | | no adduct | 782.6056 | 3 |
| | | no adduct | 786.6074 | 3 |
| | | no adduct | 790.6093 | 3 |
| | | no adduct | 794.6111 | 3 |
| | | no adduct | 798.6129 | 3 |

**Supplementary Table S12: Mass list tables for precursor ion scan (tSIM) between 800 - 900 *m/z* for MAP-MS GPF-DIA**

| Compound | Formula | Adduct | Center mass (*m/z*) | z | Isolation window (*m/z*) |
|---|---|---|---|---|---|
| Boxcar 1 | | no adduct | 808.1172 | 1 | 17 |
| Boxcar 2 | | no adduct | 825.1249 | 1 | 17 |
| Boxcar 3 | | no adduct | 842.1327 | 1 | 17 |
| Boxcar 4 | | no adduct | 859.1404 | 1 | 17 |
| Boxcar 5 | | no adduct | 876.1481 | 1 | 17 |
| Boxcar 6 | | no adduct | 893.1559 | 1 | 17 |

Table  S19

**Supplementary Table S13: Mass list tables for MS/MS scan (tMS2) between 800 - 900 *m/z* for MAP-MS GPF-DIA and standard GPF-DIA (fixed 4 *m/z* isolation windows)**

| Compound | Formula | Adduct | *m/z* | z |
|---|---|---|---|---|
| | | no adduct | 802.6147 | 3 |
| | | no adduct | 806.6165 | 3 |
| | | no adduct | 810.6184 | 3 |
| | | no adduct | 814.6202 | 3 |
| | | no adduct | 818.622 | 3 |
| | | no adduct | 822.6238 | 3 |
| | | no adduct | 826.6256 | 3 |
| | | no adduct | 830.6274 | 3 |
| | | no adduct | 834.6293 | 3 |
| | | no adduct | 838.6311 | 3 |
| | | no adduct | 842.6329 | 3 |
| | | no adduct | 846.6347 | 3 |
| | | no adduct | 850.6365 | 3 |

| | | | | |
|---|---|---|---|---|
| | | no adduct | 854.6384 | 3 |
| | | no adduct | 858.6402 | 3 |
| | | no adduct | 862.642 | 3 |
| | | no adduct | 866.6438 | 3 |
| | | no adduct | 870.6456 | 3 |
| | | no adduct | 874.6475 | 3 |
| | | no adduct | 878.6493 | 3 |
| | | no adduct | 882.6511 | 3 |
| | | no adduct | 886.6529 | 3 |
| | | no adduct | 890.6547 | 3 |
| | | no adduct | 894.6566 | 3 |
| | | no adduct | 898.6584 | 3 |

**Supplementary Table S14: Mass list tables for precursor ion scan (tSIM) between 900 - 1000 *m/z* for MAP-MS GPF-DIA**

| Compound | Formula | Adduct | Center mass (*m/z*) | z | Isolation window (*m/z*) |
|---|---|---|---|---|---|
| Boxcar 1 | | no adduct | 908.1627 | 1 | 17 |
| Boxcar 2 | | no adduct | 925.1704 | 1 | 17 |
| Boxcar 3 | | no adduct | 942.1781 | 1 | 17 |
| Boxcar 4 | | no adduct | 959.1859 | 1 | 17 |
| Boxcar 5 | | no adduct | 976.1936 | 1 | 17 |
| Boxcar 6 | | no adduct | 993.2013 | 1 | 17 |

**Supplementary Table S15: Mass list tables for MS/MS scan (tMS2) between 900 - 1000 *m/z* for MAP-MS GPF-DIA and standard GPF-DIA (fixed 4 *m/z* isolation windows)**

| Compound | Formula | Adduct | *m/z* | z |
|---|---|---|---|---|
| | | no adduct | 902.6602 | 3 |
| | | no adduct | 906.662 | 3 |
| | | no adduct | 910.6638 | 3 |
| | | no adduct | 914.6657 | 3 |
| | | no adduct | 918.6675 | 3 |
| | | no adduct | 922.6693 | 3 |
| | | no adduct | 926.6711 | 3 |
| | | no adduct | 930.6729 | 3 |
| | | no adduct | 934.6747 | 3 |
| | | no adduct | 938.6766 | 3 |
| | | no adduct | 942.6783 | 3 |
| | | no adduct | 946.6802 | 3 |
| | | no adduct | 950.682 | 3 |
| | | no adduct | 954.6838 | 3 |
| | | no adduct | 958.6857 | 3 |
| | | no adduct | 962.6875 | 3 |
| | | no adduct | 966.6893 | 3 |
| | | no adduct | 970.6911 | 3 |
| | | no adduct | 974.6929 | 3 |
| | | no adduct | 978.6948 | 3 |
| | | no adduct | 982.6965 | 3 |
| | | no adduct | 986.6984 | 3 |
| | | no adduct | 990.7002 | 3 |
| | | no adduct | 994.702 | 3 |
| | | no adduct | 998.7039 | 3 |

**Supplementary Table S16: Mass list tables for precursor ion scan (SIM) for Boxcar acquisition method.**

| Compound | *m/z* | z | MSX ID | Isolation window (*m/z*) |
|---|---|---|---|---|
| BoxCar 1-1 | 408.9538 | 1 | 1 | 17 |
| BoxCar 1-2 | 425.9435 | 1 | 2 | 17 |
| BoxCar 1-3 | 441.9508 | 1 | 3 | 15 |
| BoxCar 1-4 | 456.9576 | 1 | 4 | 15 |
| BoxCar 1-5 | 471.4642 | 1 | 5 | 14 |
| BoxCar 1-6 | 484.9703 | 1 | 6 | 13 |
| BoxCar 2-1 | 498.4765 | 1 | 1 | 14 |
| BoxCar 2-2 | 511.9826 | 1 | 2 | 13 |
| BoxCar 2-3 | 524.9885 | 1 | 3 | 13 |
| BoxCar 2-4 | 537.9944 | 1 | 4 | 13 |
| BoxCar 2-5 | 551.0004 | 1 | 5 | 13 |
| BoxCar 2-6 | 563.506 | 1 | 6 | 12 |
| BoxCar 3-1 | 576.0117 | 1 | 1 | 13 |
| BoxCar 3-2 | 589.0176 | 1 | 2 | 13 |
| BoxCar 3-3 | 602.0236 | 1 | 3 | 13 |
| BoxCar 3-4 | 615.0294 | 1 | 4 | 13 |
| BoxCar 3-5 | 628.0354 | 1 | 5 | 13 |
| BoxCar 3-6 | 641.0413 | 1 | 6 | 13 |
| BoxCar 4-1 | 654.5474 | 1 | 1 | 14 |
| BoxCar 4-2 | 668.0536 | 1 | 2 | 13 |
| BoxCar 4-3 | 681.5597 | 1 | 3 | 14 |
| BoxCar 4-4 | 695.566 | 1 | 4 | 14 |
| BoxCar 4-5 | 709.5724 | 1 | 5 | 14 |
| BoxCar 4-6 | 724.079 | 1 | 6 | 15 |
| BoxCar 5-1 | 739.5861 | 1 | 1 | 16 |

| | | | | |
|---|---|---|---|---|
| BoxCar 5-2 | 756.0936 | 1 | 2 | 17 |
| BoxCar 5-3 | 773.1013 | 1 | 3 | 17 |
| BoxCar 5-4 | 790.6093 | 1 | 4 | 18 |
| Boxcar 5-5 | 808.6174 | 1 | 5 | 18 |
| Boxcar 5-6 | 827.6261 | 1 | 6 | 20 |
| BoxCar 6-1 | 848.1354 | 1 | 1 | 21 |
| BoxCar 6-2 | 870.1454 | 1 | 2 | 23 |
| BoxCar 6-3 | 894.1563 | 1 | 3 | 25 |
| BoxCar 6-4 | 920.6683 | 1 | 4 | 28 |
| BoxCar 6-5 | 949.6815 | 1 | 5 | 30 |
| BoxCar 6-6 | 982.6965 | 1 | 6 | 36 |

**Supplementary Table S17: Isolation windows for alternate MAP-MS x3 method**

| Compound | Formula | Adduct | Precursor (m/z) | Precursor Charge (z) | MSX ID | Isolation window (*m/z*) |
|---|---|---|---|---|---|---|
| Boxcar1 | | (no adduct) | 484.9703 | 1 | 1 | 169 |
| Boxcar2 | | (no adduct) | 650.5455 | 1 | 1 | 162 |
| Boxcar3 | | (no adduct) | 866.1436 | 1 | 1 | 269.1 |

**Supplementary Table S18: Isolation windows for alternate MAP-MS x12 method**

| Compound | Formula | Adduct | Precursor (m/z) | Precursor Charge (z) | MSX ID | Isolation window (*m/z*) |
|---|---|---|---|---|---|---|
| Boxcar1 | | (no adduct) | 424.9431 | 1 | 1 | 49 |
| Boxcar2 | | (no adduct) | 470.4638 | 1 | 1 | 42 |
| Boxcar3 | | (no adduct) | 511.4824 | 1 | 1 | 40 |
| Boxcar4 | | (no adduct) | 550.5001 | 1 | 1 | 38 |
| Boxcar5 | | (no adduct) | 589.0176 | 1 | 1 | 39 |
| Boxcar6 | | (no adduct) | 628.0354 | 1 | 1 | 39 |
| Boxcar7 | | (no adduct) | 668.0536 | 1 | 1 | 41 |
| Boxcar8 | | (no adduct) | 710.0726 | 1 | 1 | 43 |
| Boxcar9 | | (no adduct) | 756.5938 | 1 | 1 | 50 |
| Boxcar10 | | (no adduct) | 809.6179 | 1 | 1 | 56 |
| Boxcar11 | | (no adduct) | 872.1463 | 1 | 1 | 69 |
| Boxcar12 | | (no adduct) | 953.6834 | 1 | 1 | 94 |

**Supplementary Table S19: Isolation windows for alternate MAP-MS x20 method**

| Compound | Formula | Adduct | Precursor (m/z) | Precursor Charge (z) | MSX ID | Isolation window (*m/z*) |
|---|---|---|---|---|---|---|
| Boxcar1 | | (no adduct) | 415.9389 | 1 | 1 | 31 |
| Boxcar2 | | (no adduct) | 444.9522 | 1 | 1 | 27 |
| Boxcar3 | | (no adduct) | 470.964 | 1 | 1 | 25 |
| Boxcar4 | | (no adduct) | 495.9753 | 1 | 1 | 25 |
| Boxcar5 | | (no adduct) | 519.9862 | 1 | 1 | 23 |
| Boxcar6 | | (no adduct) | 542.9967 | 1 | 1 | 23 |
| Boxcar7 | | (no adduct) | 566.0072 | 1 | 1 | 23 |
| Boxcar8 | | (no adduct) | 589.0176 | 1 | 1 | 23 |
| Boxcar9 | | (no adduct) | 612.0281 | 1 | 1 | 23 |
| Boxcar10 | | (no adduct) | 635.5388 | 1 | 1 | 24 |
| Boxcar11 | | (no adduct) | 659.5497 | 1 | 1 | 24 |
| Boxcar12 | | (no adduct) | 684.0608 | 1 | 1 | 25 |
| Boxcar13 | | (no adduct) | 709.5724 | 1 | 1 | 26 |
| Boxcar14 | | (no adduct) | 737.0849 | 1 | 1 | 29 |
| Boxcar15 | | (no adduct) | 766.5983 | 1 | 1 | 30 |
| Boxcar16 | | (no adduct) | 798.1127 | 1 | 1 | 33 |
| Boxcar17 | | (no adduct) | 832.6284 | 1 | 1 | 36 |
| Boxcar18 | | (no adduct) | 871.1459 | 1 | 1 | 41 |
| Boxcar19 | | (no adduct) | 916.1664 | 1 | 1 | 49 |

| | | | | | | |
|---|---|---|---|---|---|---|
| Boxcar20 | | (no adduct) | 970.6912 | 1 | 1 | 60 |

**Supplementary Table S20: Number of identified peptides and their corresponding charge states detected for each acquisition method.**

| Method | AGC target | Single charge | 2 charge states | 3 charge states |
|---|---|---|---|---|
| MAP-MS | 1e+06 | 69,807 | 9,854 | 73 |
| Standard | 1e+06 | 68,177 | 7,026 | 55 |
| Standard | 3e+06 | 69,034 | 6,993 | 41 |
